# Lamp1 mediates lipid transport, but is dispensable for autophagy in *Drosophila*

**DOI:** 10.1101/2021.03.03.432938

**Authors:** Norin Chaudhry, Margaux Sica, Satya Surabhi, David Sanchez Hernandez, Ana Mesquita, Adem Selimovic, Ayesha Riaz, Hua Bai, Gustavo C. Macintosh, Andreas Jenny

**Affiliations:** Roy J. Carver Dept. of Biochemistry, Biophysics and Molecular Biology, Iowa State University, Ames, IA, USA; Department of Developmental and Molecular Biology, New York, NY, USA; Department of Genetics, Albert Einstein College of Medicine, New York, NY, USA; Department of Genetics, Development and Cell Biology, Iowa State University, Ames, IA, USA

**Keywords:** LAMP proteins, autophagy, *Drosophila*, lipid transport, lysosome

## Abstract

The endolysosomal system not only is an integral part of the cellular catabolic machinery that processes and recycles nutrients for synthesis of biomaterials, but also acts as signaling hub to sense and coordinate the energy state of cells with growth and differentiation. Lysosomal dysfunction adversely influences vesicular transport-dependent macromolecular degradation and thus causes serious problems for human health. In mammalian cells, loss of the lysosome associated membrane proteins LAMP1/2 strongly impacts autophagy and cholesterol trafficking. Here we show that the previously uncharacterized *Drosophila* Lamp1 is a *bona fide* homolog of vertebrate LAMP1/2. Surprisingly and in contrast to *Lamp1/2* double mutant mice, *Drosophila* Lamp1 is not required for viability or autophagy, suggesting that autophagy defects in *Lamp1/2* mutants may have indirect causes. However, Lamp1 deficiency results in an expansion of the acidic compartment in flies. Furthermore, we find that *Lamp1* mutant larvae have defects in lipid metabolism as they show elevated levels of sterols and diacylglycerols (DAGs). Since DAGs are the main lipid species used for transport though the hemolymph (blood) in insects, our results indicate broader functions of Lamp1 in lipid transport. Our findings make *Drosophila* an ideal model to study the role of LAMP proteins in lipid assimilation without the confounding effects of their storage and without interfering with autophagic processes.

## Introduction

Endocytosis, phagocytosis, and autophagy funnel proteins and lipids into lysosomes where they are degraded by hydrolases in their acidic environment [1-4]. Lysosomes, originally identified by DeDuve in the 1950s, were thus traditionally considered catabolic organelles degrading biomass to recycle metabolic building blocks for biosynthetic processes. Only more recently, the lysosomal surface has also been recognized as signaling hub regulating nutrient signaling and lysosomal biogenesis via recruitment of the nutrient sensor mTORC1 (mechanistic target of rapamycin) or the master regulator of lysosomal biosynthesis TFEB (transcription factor EB; reviewed in [5-8]). Lysosomal dysfunction affects membrane trafficking and repair on the one hand, and metabolism and signaling on the other [5-8] and thus has wide-ranging consequences for human health. Lysosomal storage diseases are caused by mutations in about 70 genes encoding hydrolases, lysosomal membrane proteins (LMPs), and transport proteins [8] and include Pompe disease caused by glycogen accumulation in lysosomes [9] and Nieman Pick Type C disorder resulting in lysosomal lipid, particularly cholesterol, accumulation [10]. Additionally, degradation of cytoplasmic proteins and organelles via autophagy is critically dependent on lysosomal function, and defective lysosomal proteolysis and autophagosomal-lysosomal fusion contribute to neurodegenerative diseases including Parkinson, Alzheimer’s and Huntington disease [3,11,12].

The LAMP (lysosome associated membrane protein) family of proteins are type 1 transmembrane domain proteins characterized by luminal LAMP domains stabilized by two S-S bridges (Fig. 1A) [13,14] that are followed by a transmembrane domain and short C-terminus facing the cytosol and containing a YXXΦ-type endosomal sorting signal (Fig. 1A) [15-17]. In vertebrates, LAMP1 and LAMP2 make up ∼50% of proteins in the lysosomal membrane [18,19]. Their luminal domains are heavily N- and O-glycosylated and form a glycocalyx that is hypothesized to protect the lysosomal membrane from the acidic hydrolases [18]. While *Lamp1* mutant mice are mostly normal [20], 50% of *Lamp2* mutants die within weeks of birth, and surviving animals show cardiomyopathy and an increase in autophagic vacuoles (AVs) in many tissues including the liver [21,22]. Importantly, human patients with mutations in *LAMP2* have Danon disease, characterized by similar phenotypes [23,24]. *LAMP2* encodes three splice-isoforms that differ in their TM region and C-termini. Lack of the 2B isoform is sufficient to cause defects in macroautophagy (MA), the form of autophagy dependent on the formation of autophagosomes which engulf cytoplasmic content [21,22,24-26]. Arguably, the best characterized isoform is LAMP2A, which functions as substrate translocation channel in chaperone mediated autophagy (CMA), a form of autophagy specific for soluble cytoplasmic proteins containing a KFERQ-targeting motif. Lack of LAMP2A leads to premature aging-like phenotypes and enhances neurodegeneration [27-29]. *Lamp1/2* double-mutant mice are embryonic lethal and double-mutant mouse embryonic fibroblasts (MEF) show a strong accumulation of autophagic vacuoles (AV) and unesterified cholesterol in late endosomes and lysosomes [2,18,25,30]. Furthermore, *Lamp2* and *Lamp1/2* double mutant MEFs and hiPSC-derived cardiomyocytes from Danon disease patients revealed a block of MA flux [25,31]. However, the cause for the accumulation of AVs and the block of MA is not understood and could be indirect [21,25].

**Figure 1.**
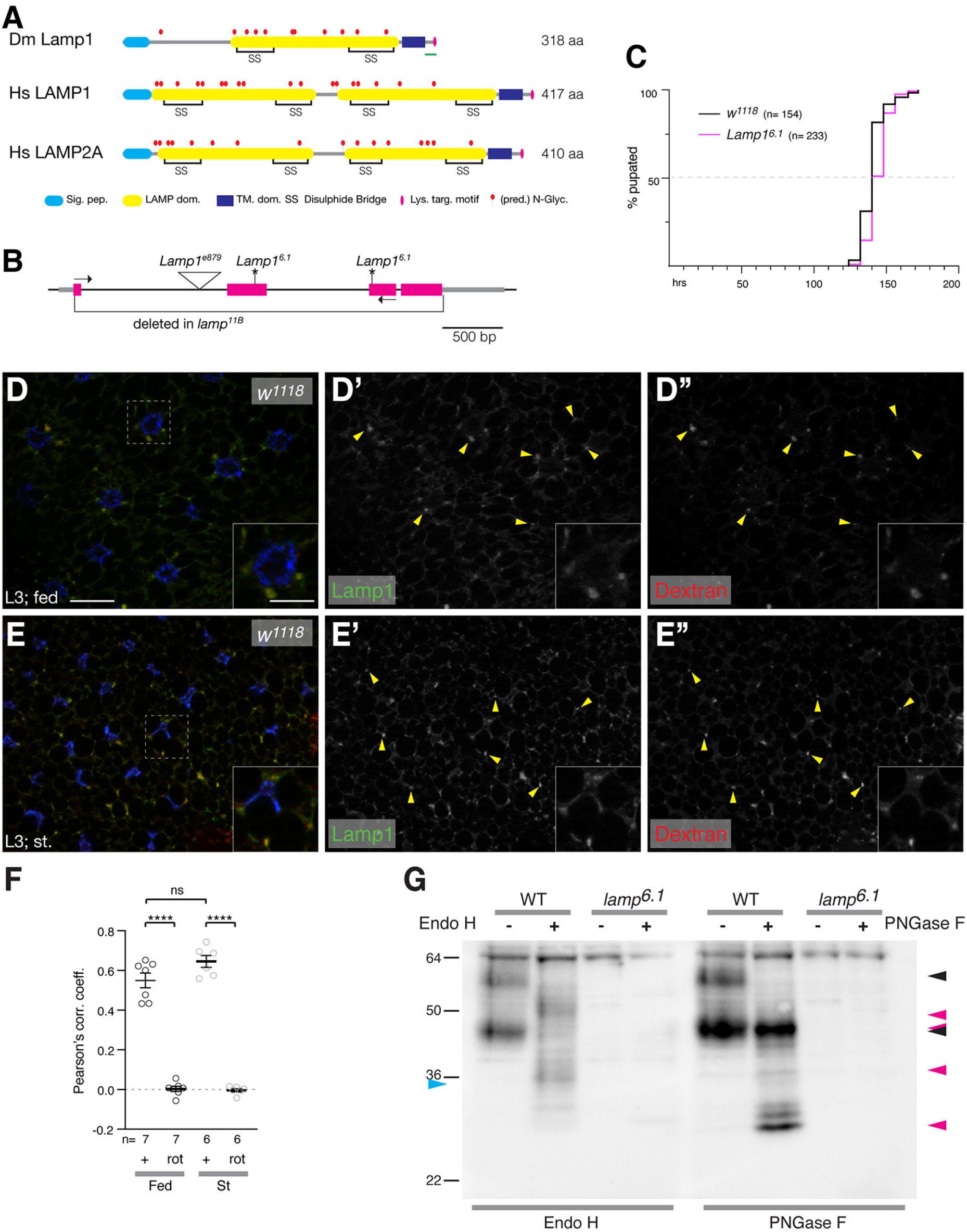
*Lamp1* mutants are viable and have no developmental delay. (A) Schematic comparing *Drosophila* Lamp1 with human LAMP1 and LAMP2A. Note that Lamp1 has only one luminal LAMP domain. The green bar indicates the peptide region used to raise the Lamp1 antibody. (B) Schematic of *Lamp1* locus with mutant alleles indicated. UTRs are in grey, coding sequence in magenta. *Lamp1*^*6*.*1*^ contains two frame shifts (*) at the positions of both gRNAs used to induce it. Arrows depict location of RT-PCR primers. (C) Quantification of pupation timing shows that *Lamp1*^*6*.*1*^ mutants have no developmental delay. n indicates total number of pupae scored. (D, E) In 3^rd^ instar larval fat body under fed (C) and starved conditions (D), Lamp1 colocalizes with TRITC-Dextran-labeled endolysosomes (examples marked by yellow arrow heads). Single channel images show Lamp1 (D’, E’) and Dextran (D”, E”), respectively. Insets show enlaged section indicated by dotted squares. Nuclei are in blue. Scale bars are 20 µm (10 µm in insets). (F) Pearson’s correlation coefficient for the colocalization of Lamp1 with TRITC-Dextran under indicated conditions. As control, one channel was rotated (rot) by 180°. One-way ANOVAs (Tukey correction) P <0.0001; ****, P <0.0001; ns, not significant. (G) Western blot of adult head lysates untreated (-) or treated with indicated glycosidase (+) form *w*^*1118*^ (WT) and *Lamp1*^*6*.*1*^ mutant flies. Blue arrowhead indicates predicted molecular mass, black arrowheads untreated Lamp1, magenta arrowheads deglycosylated forms.

In contrast to the two partially redundant mouse *Lamp1/2* genes, the *Drosophila melanogaster* genome contains a single annotated *Lamp1* gene evolutionarily related to vertebrate *LAMP1/2* that remains uncharacterized [32,33]. To our surprise, we find that *Drosophila Lamp1* null mutants are homozygous viable and show no delay during development, suggesting normal steroid hormone signaling. Furthermore, *Lamp1* mutants have no defect in macroautophagy or endosomal microautophagy (eMI), suggesting that LAMP proteins are not *per se* required for autophagy. However, we find a strong increase of acidic vesicles under basal and starvation conditions in the fat body that has functions akin to the mammalian liver. Furthermore, lipid analyses show that while levels of neutral fats (triacylglycerols; TAG) are unaffected in larvae, levels of sterols and medium chain diacylglycerols (DAG) are increased, denoting changes in lipid transport. Our results thus indicate that the roles of LAMP proteins in autophagy and in lipid homeostasis are likely independent processes.

## Results

### Drosophila Lamp1 localizes to endolysosomes and is dispensable for viability

*Drosophila Lamp1* (CG3305) encodes a protein of 318 aa with a N-terminal signal peptide followed by a single LAMP domain with four conserved cysteines, which is 21% and 18% identical to the membrane proximal and 16% and 17% identical to the distal LAMP domain of human LAMP1 and LAMP2, respectively (Clustal 2.1) [34]. These characteristics suggest that it is the homolog of the mammalian *LAMP1/2* genes (Fig. 1A) [32]. To determine the subcellular localization of Lamp1, we stained 3^rd^ instar larval fat bodies (FB) for Lamp1 with an antibody that we generated against its C-terminal peptide (Fig. 1A). Lamp1 localizes in a punctate pattern under fed and starved conditions (Fig. S1 E, G), a pattern that is specific, as no staining is found in *Lamp1*^*6*.*1*^ mutants (Fig. S1F, H; see below for mutant). To determine the identity of Lamp1-associated organelles, we labeled late endosomes and lysosomes of 3^rd^ instar larval FB using fluid phase endocytosis of fluorescent dextran (90 min chase period) [35,36] followed by staining for Lamp1. Under fed and starved conditions Lamp1 strongly co-localizes with fluorescent dextran (Fig. 1D and E, respectively), also reflected in the quantification of Pearson’s correlation coefficients of 0.6 and 0.7, respectively (Fig. 1F) [37].

To address the function of Lamp1, we generated mutants using CRISPR-Cas9 [38] and obtained two independent alleles (Fig. 1B). *Lamp1*^*6*.*1*^ contains frameshifts in exons two and three, well upstream of the TM domain, possibly encoding the first 90 aa of Lamp1, while in *Lamp1*^*11B*^, all but the first 8 aa are deleted (Figs. 1B and S1A). Additionally, the PiggyBac insertion *Lamp1*^*e879*^, inserted in the first intron of *Lamp1*, is an RNA null allele, as RT-PCR showed absence of any mRNA (Fig. S1B). Unexpectedly, all three *Lamp1* alleles are homozygous viable and fertile and adults do not show an externally visible phenotype. In addition, *Lamp1*^*6*.*1*^ mutants show no developmental delay, as they take a median of 140 h to pupation similar to WT controls (Fig. 1C).

Lamp1 protein migrates at an apparent Mw of 45 kDa with additional larger bands of up to ∼ 60 kDa in lysates of adult heads (Fig. S1C) or 3^rd^ instar larvae (Fig. S1D), thus considerably larger than its predicted Mw of 34.8 kDa, consistent with 11 predicted N-glycosylation sites upstream of a transmembrane domain (Fig. 1A). Importantly, these bands are specific to Lamp1, as they are absent from lysates of *Lamp1*^*6*.*1*^ and *Lamp1*^*11B*^ mutants (Fig. S1C,D), also suggesting that all *Lamp1* alleles are likely null alleles. Upon treatment of adult head lysates with Endo H, which removes mannose rich oligosaccharides from proteins [39,40], or PNGase F that removes all N-linked sugars [40], the apparent Mw of Lamp1 specific bands shifts towards smaller protein species at the expense of the largest ones, an effect that is stronger upon PNGase F treatment (Fig. 1G). The late endosome and lysosomal localization, and N-glycosylation status of Lamp1 are consistent with it being a *bona fide* homolog of mammalian LAMP1/2.

### Expansion of the acidic endolysosomal compartment in Lamp1 mutants

To assess changes in the endolysosomal compartment, we stained larval FB *ex vivo* with lysotracker LTR, that is retained in acidic structures upon protonation and has been used to monitor starvation induced MA as barely any Lysotracker positive structures are found under fed conditions [41]. Compared to control (Fig. 2A), FB of well-fed larvae of all three *Lamp1* alleles show very pronounced increase in acidic structures (Fig. 2 B,C and Fig. S2; quantified in Fig. 2I). Importantly, this phenotype is rescued by a duplication that includes the *Lamp1* locus (*Lamp1*^*Dp*^; Fig. 2D, quantified in I) and the phenotype is not restricted to larvae, but also found in FB of adult *Lamp*^*e879*^ mutants (Fig. 2G,H; quantified in 2J). A similar, albeit smaller expansion of acidic structures is found upon starvation (Figs. 2E, F and S2D-F; quantified in 2I). The increase in LTR staining is not due to defects in the regulation of a starvation response in *Lamp1* mutants, as the mRNA of *Amyrel*, the gene encoding an α-Amylase-like protein, known to be strongly induced by starvation [42,43], is low in fed larvae and normally upregulated by starvation to similar levels in WT and *Lamp1* mutants (Fig. 2K).

**Figure 2.**
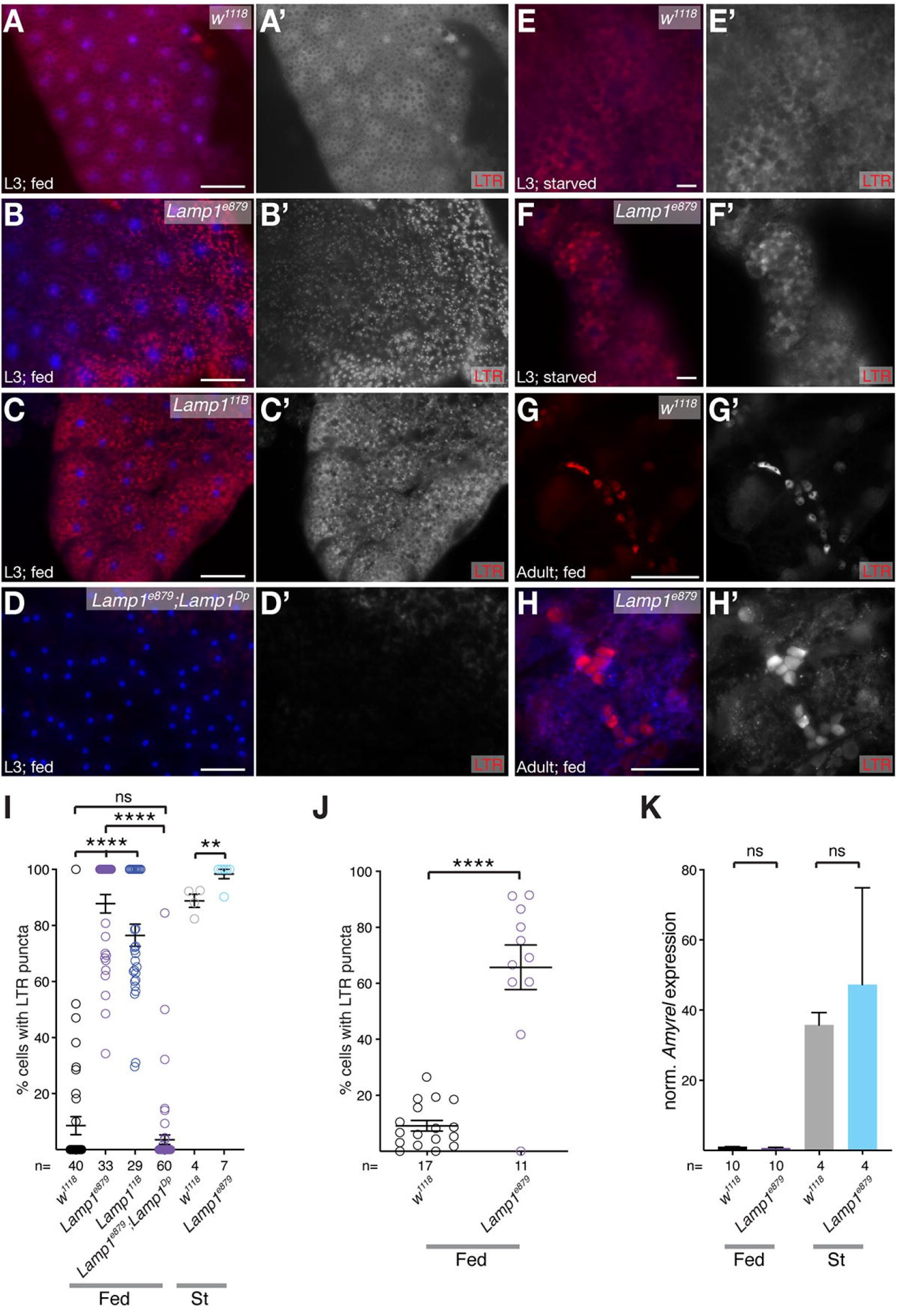
*Lamp1* mutants have strongly increased acidic structures. Compared to *w*^*1118*^ (A), fed 3^rd^ instar FB of *Lamp1*^*e879*^ (B) *Lamp1*^*11B*^ (C) show a strong increase of LTR staining, a phenotype that is rescued by a duplication including Lamp1 (D; quantified in I). (E, F) LTR staining is also increased in starved 3^rd^ instar FB of *Lamp1*^*e879*^ mutants (compare E, F; quantified in I). (G, H) A similar increase in the acidic compartment is seen in adult fat body of *Lamp1*^*e879*^ (compare G, H; quantified in J). Nuclei are in blue; greyscale images show LTR channel; scale bars: 20 µm. (I, J) Quantification LTR staining in larval (I) and adult (J) fat body of indicated genotypes. One-way ANOVA (Tukey correction) for Fed in (I) P <0.0001; T-tests (J); **P <0.01, ****, P <0.0001; ns, not significant. (K) *Amyrel* is normally induced upon starvation in *Lamp1*^*e879*^ mutant larvae (qPCR normalized to RpL32). Two-tailed T-test.

To test if the LTR phenotype correlated with increased lysosomal activity, we first stained FB of 3^rd^ instar larvae with MagicRed, a Cathepsin B substrate that becomes fluorescent in active, acidic lysosomes [44,45]. Under basal, fed conditions and in contrast to increased LTR staining, *Lamp1* mutants show no increase of Cathepsin B activity, suggestive of unchanged lysosomal activation (Fig. 3A-C; compare *w*^*1118*^ with *Lamp1*^*6*.*1*^ (B) and *Lamp1*^*11B*^ (C); quantified in 3G). This is also supported by normal Acid Phosphatase activity in lysosome enriched fractions of *Lamp1* mutant larval lysates (Fig. 3H), a common marker of lysosomal activity [46,47]. However, although quite variable, Cathepsin B activity is higher in *Lamp1* mutants under starved conditions (Fig. 3D-F; quantified in 3G), suggesting that lysosomal activation can be abnormal under certain stress conditions.

**Figure 3.**
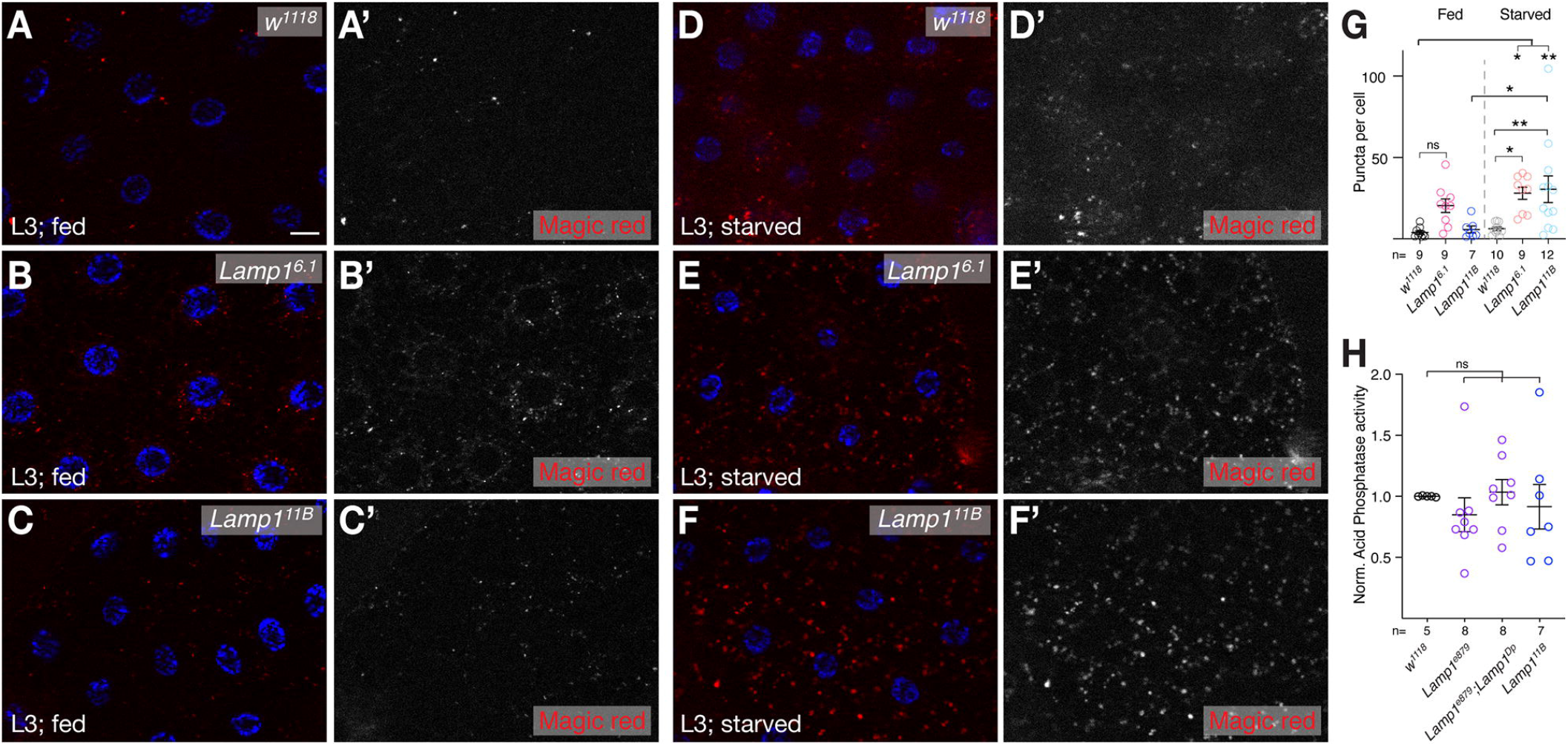
Lysosomal activity. (A-C) Cathepsin B activity as measured by Magic Red assay is not significantly increased in FB of fed 3^rd^ instar larvae (compare *w*^*1118*^ in A with *Lamp1*^*6*.*1*^ (B) and *Lamp1*^*11B*^ (C); quantified in G). In contrast, under starved conditions (D-F), *Lamp1*^*6*.*1*^ (E) and *Lamp1*^*11B*^ (F) have increased lysosomal activity (compare to *w*^*1118*^ in D; quantified in G). Nuclei are in blue; greyscale images show Magic Red channel; scale bar: 20 µm. (G) Quantification of Magic Red puncta in FB of indicated genotypes. One-way ANOVA (Tukey correction) P <0.001. (H) Normalized Acid Phosphatase activity of lysates enriched for lysosomes of fed 3^rd^ instar larvae of indicated genotypes. One-way ANOVA (Dunnett correction). *, P <0.05; **P <0.01, ns, not significant.

### Lamp1 is dispensable for autophagy

The expanded acidic compartment could be suggestive of a MA defect. MA is well characterized in 3^rd^ instar fat body [36,48,49] and autophagic flux is commonly assessed using a tandem-tagged GFP-mCherry-Atg8a reporter which fluoresces green and red in autophagosomes, but red only in autolysosomes due to GFP fluorescence being quenched by their acidic pH [50]. However, neither under fed (Fig. 4 B; quantified in 4E) nor starved conditions (Fig. 4D; quantified in 4E) do *Lamp1*^*6*.*1*^/ *Lamp1*^*11B*^ transheterozygous or *Lamp1*^*6*.*1*^ homozygous mutants (quantified in Fig. 4E) show a difference in APGs (green puncta) nor total autophagic structures (red puncta) compared to controls (Fig. 4A, B; quantified in 4E). Importantly, we find a robust, normal stimulation of MA upon starvation as seen in the strong increase of autolysosomes and a tendency towards an increase in APGs (Fig. 4C, D; quantified in Fig. 4E). Prior to fusion of APGs with lysosomes, APGs recruit the SNAREs Syx17 (STX17 in vertebrates) and Snap29 as part of the fusion process [51,52]. Consistent with functional starvation induced MA, Snap29 is normally recruited to autophagosomes marked by mCherry-Atg8a of 3^rd^ instar FB cells of *Lamp1*^*6*.*1*^ homozygous mutants (quantified in Fig. 4H) or *Lamp1*^*6*.*1*^/ *Lamp1*^*11B*^ transheterozygotes (Fig. 4F,G; quantification in 4H). In addition, there is no change in the expression of the core MA genes *Atg5* and *Atg8a* in adult flies (Fig. 4E). Our data thus suggest, that, in contrast to humans and mice [21-24], there is no basal or starvation induced MA defect in the absence of Lamp1 in *Drosophila*.

**Figure 4.**
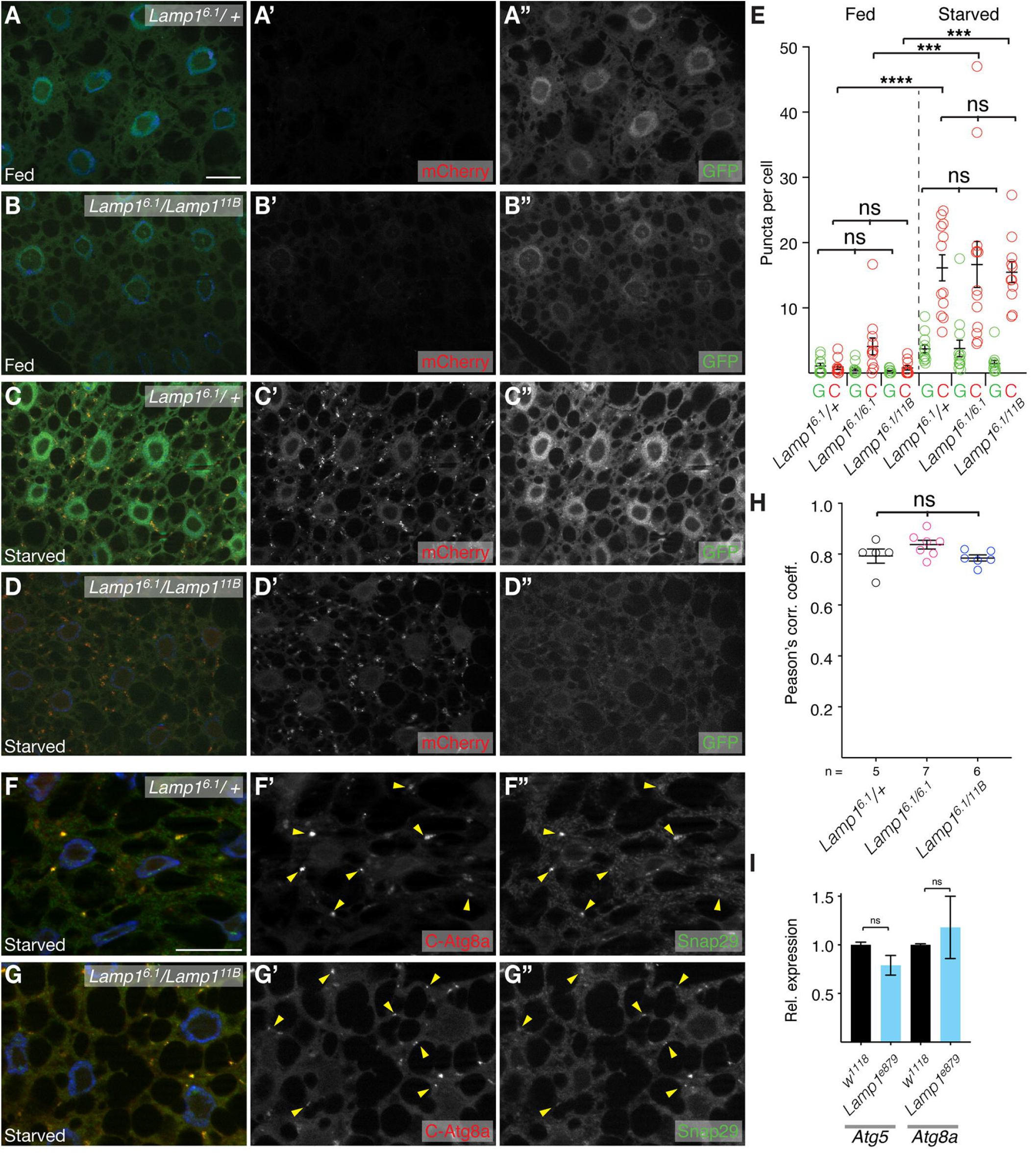
Autophagic flux is normal in FB of *Lamp1* mutants. (A,B) Under basal conditions, MA in 3^rd^ instar FB is low and indistinguishable between *Lamp1*^*6*.*1*^ heterozygous control (A), *Lamp1*^*6*.*1*^/ *Lamp1*^*11B*^ (B), and homozygous *Lamp1*^*6*.*1*^ (quantified in E) mutants. (C, D) MA is normally induced by starvation in *Lamp1*^*6*.*1*^ heterozygotes (C), *Lamp1*^*6*.*1*^/ *Lamp1*^*11B*^ (D), and homozygous *Lamp1*^*6*.*1*^ (quantified in E). APGs are labeled with GFP and mCherry, while GFP fluorescence is quenched by the acidic pH in autolysosomes, which are thus labeled in red only. (E) Quantification of GFP puncta (APGs) and cherry puncta (APGs and APGLs) shows normal MA flux in *Lamp1* mutants with no difference to controls. One-way ANOVA (Tukey correction) P< 0.0001. (F, G) Snap29 (green) recruitment to APGs (marked by mCherry-Atg8a; C-Atg8a; red) is normal in FB of *Lamp1*^*6*.*1*^/ *Lamp1*^*11B*^ transheterozygotes (G) and *Lamp1*^*6*.*1*^ mutants (quantified in H; compare to heterozygotes in F; examples of APGs are marked by yellow arrowheads). (H) Pearson’s correlation coefficient of colocalization of Snap29 with mCherry-Atg8a of indicated genotypes. One-way ANOVA; Dunnett correction) ns. Nuclei are in blue; greyscale images show indicated channels; scale bars: 20 µm. (I) *Lamp1*^*e879*^ mutant adults have normal expression of the essential MA genes *Atg5* and *Atg8* (fed; normalized to *RpL32*). Two-tailed T-tests. ***P <0.001, ****, P <0.0001; ns, not significant.

In mammals, in addition to MA, CMA and eMI also contribute to autophagic protein degradation [1-4]. eMI has recently been identified in flies based on its selective requirement of Hsc70-4 and the ESCRT machinery [53,54]. On the other hand, to date, CMA has only been shown to occur in mammals and fish and other species (except for birds) lack amino acids critical for Hsc70/HSPA8 interaction in their LAMP proteins [28,55]. We therefore tested if Lamp1 is required for eMI, but found that similar to MA, there is no eMI defect in *Lamp1* mutants, as prolonged starvation normally induces eMI sensor puncta (KFERQ-PAmCherry) in *Lamp1*^*6*.*1*^ mutants (Fig. 5; compare heterozygous larval FB (A,C) with homozygous mutant tissue (B,D); quantification in 5E). Given the essential role of LAMP2A in CMA and that fact that KFERQ-PAmCherry acts as CMA sensor in vertebrates [56,57], the lack of differences in the levels of KFERQ-mCherry puncta between wild-type and *Lamp1* mutants support the idea that flies lack a CMA-like process.

**Figure 5.**
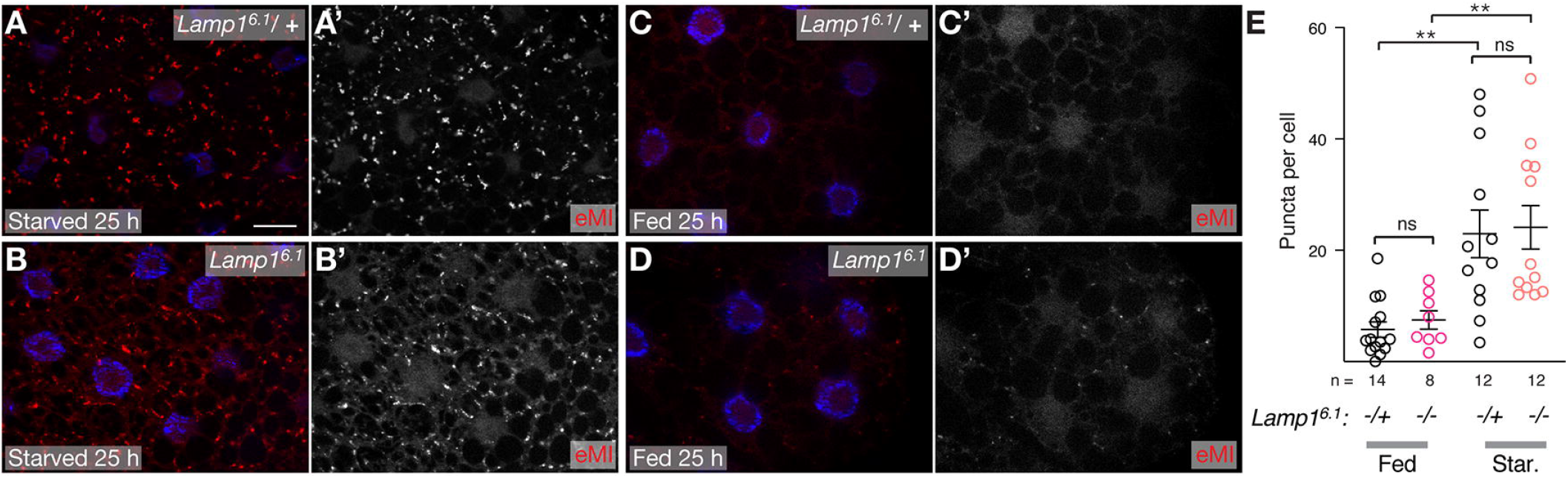
eMI induced by prolonged starvation is normal in *Lamp1*^*6*.*1*^ mutants (compare starved *Lamp1*^*6*.*1*^ heterozygous control (A) and homozygous *Lamp1*^*6*.*1*^ mutant larval FB (B) with corresponding fed tissue (C,D). Greyscale images show eMI sensor; nuclei are in blue; scale bar 20µm. (E) quantification of eMI sensor puncta per 3^rd^ instar FB cell of indicated genotypes. One-way ANOVA (Tukey correction) P <0.0001; **, P <0.01; ns, not significant.

To independently assess MA, we performed morphometric analyses of autophagic structures on TEM sections of fed and starved *Lamp1*^*6*.*1*^ mutant 3^rd^ instar FB (Fig. 6). No difference between wild-type and *Lamp1* mutant tissue was found for the density of mature autolysosomes (examples in Fig. 6A, C; quantified in Fig. 6G; see materials and methods for definition of structures) and lysosomes (examples in Fig. 6D, E; quantified in Fig. 6H). A tendency towards an increased area of APGLs was found for fed *Lamp1*^*6*.*1*^ mutant FB, while the size distribution of those structures changed towards smaller ones under starvation (Fig. 6I). Autophagosomes (AVi in e.g. [18]) were very rare in all cases. Overall, our data thus show no indication for autophagy defects in *Lamp1* mutants.

**Figure 6.**
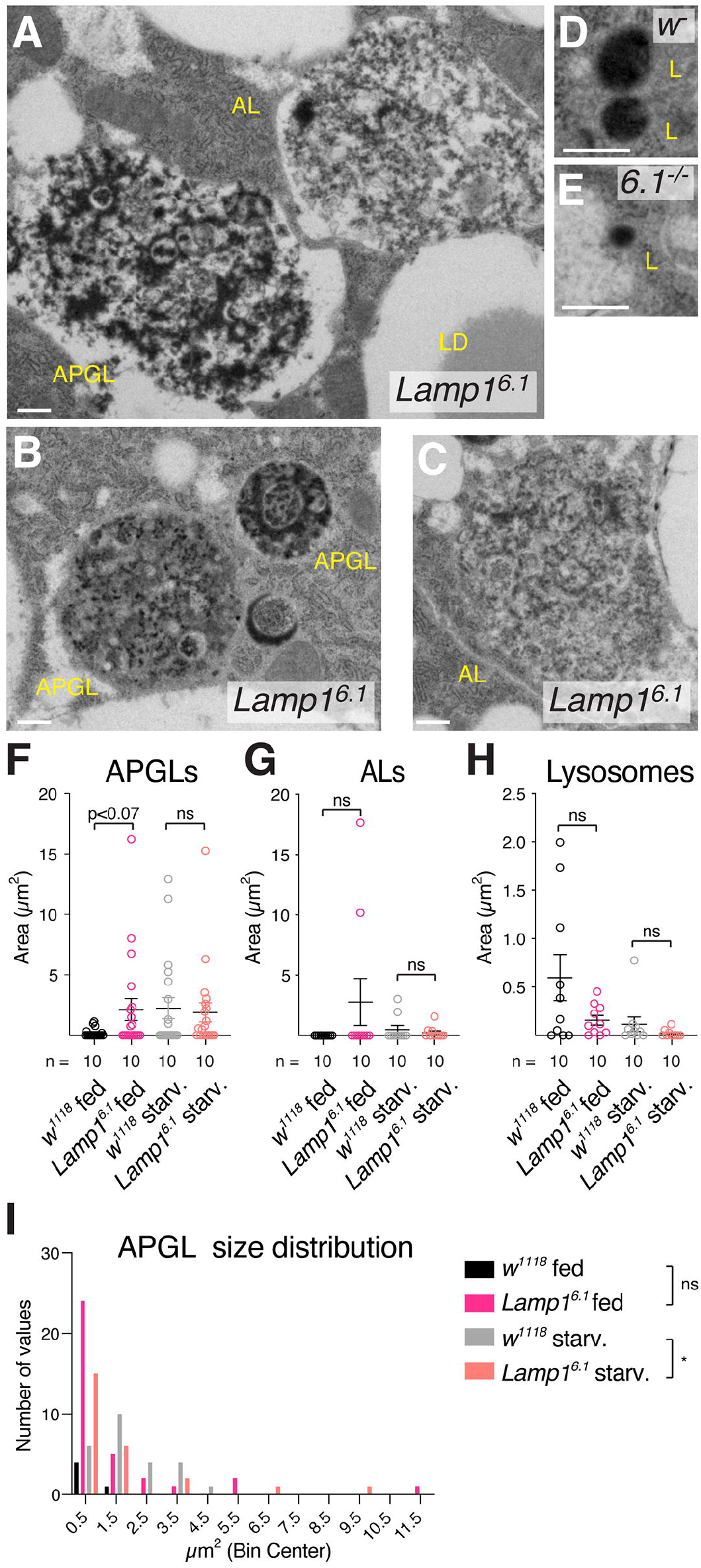
TEM analyses of *Lamp1* mutant fat body. (A-E) Examples of autolysosomal structures of *Lamp1*^*6*.*1*^ mutant (A-C) and WT (D) 3^rd^ instar fed larval fat body. APGL: autophagolysosome; AL: autolysosome; L: lysosome; LD: lipid droplet. See also methods for structure definitions. Scale bars: 0.5 µm. (F-H) Quantification of density (area per field of view) of APGLs (F), ALs (G), and lysosomes (H) under indicated conditions. Mann-Whitney tests. (I) Size distribution of APGLs under indicated conditions. Mann-Whitney and Kolmogorov-Smirnov tests. *, P <0.05; ns: not significant.

### Altered lipid content in *Lamp1* mutants

*Lamp1/2* mutant MEFs accumulate cholesterol in the endolysosomal system and it was shown that LAMP1/2 proteins bind cholesterol for storage in lysosomes [26,58]. We therefore determined whether lack of Lamp1 altered lipid profiles of mutant 3^rd^ instar larvae. *Lamp1*^*e879*^ mutants showed elevated total sterol levels, a phenotype rescued by a *Lamp1* duplication (Fig. 7A). In flies, dietary lipids are taken up by enterocytes in the gut and converted to diacylglycerols (DAG), secreted and transported in the hemolymph (blood) as lipoprotein complexes, from where they are taken up by other tissues such as FB cells that convert them to triacyclglycerols (TAGs) for storage [59]. No changes were found in triglycerides in *Lamp1*^*e879*^ mutants (Fig. 7B), consistent with no change in lipid droplet density (LD) in the fat body in our TEM analyses (Fig. S3A). In contrast, *Lamp1* mutant larvae show elevated levels of DAGs that are rescued by *Lamp1* duplication (Fig. 7C). The increase is specific for DAGs with medium chain fatty acid tails (Fig. 7C), characteristic of DAGs associated with lipoproteins, which suggests that lipid transport or mobilization is affected. No changes were found on glycogen density by TEM (Fig. S3C).

**Figure 7.**
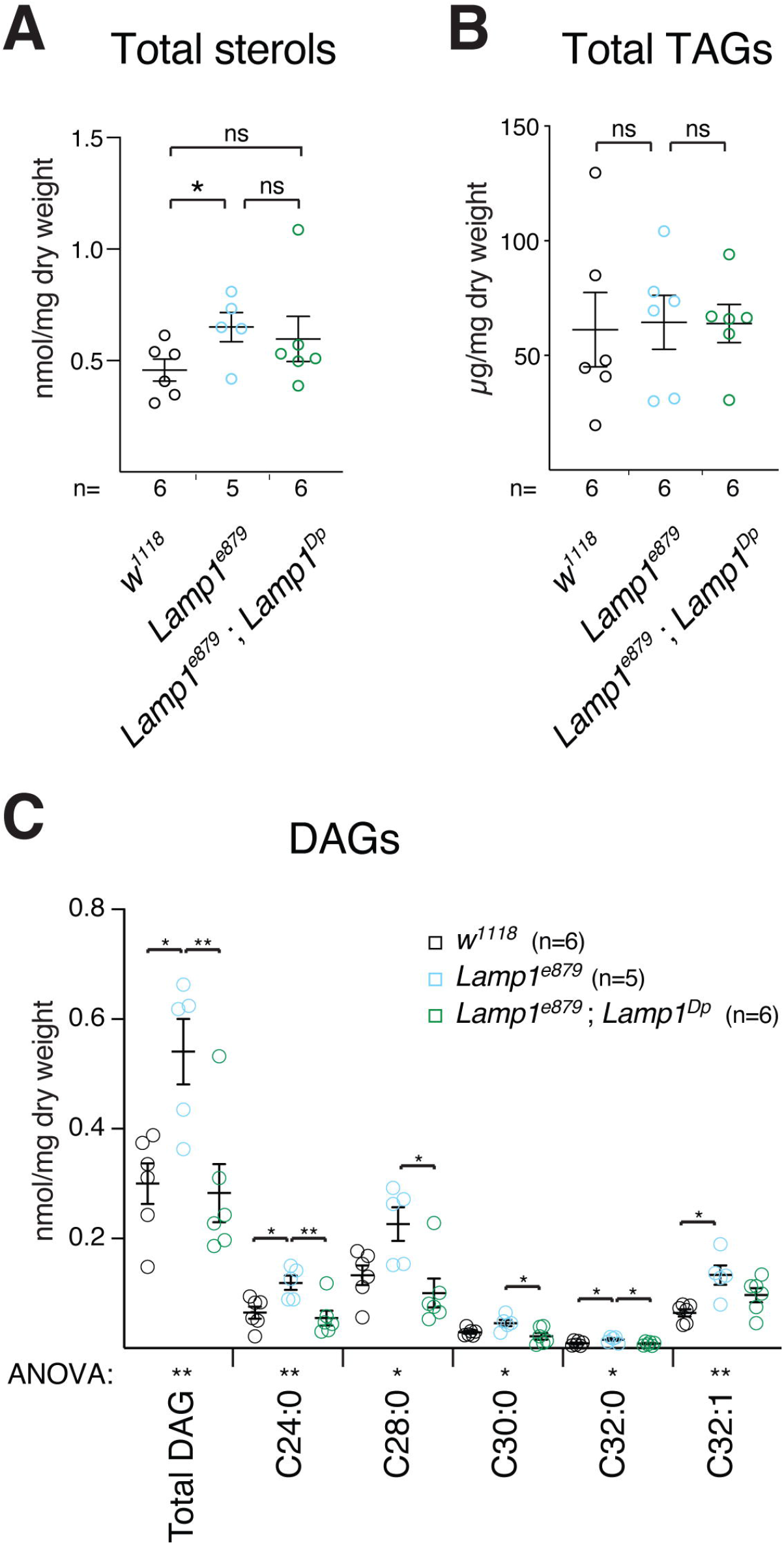
*Lamp1* mutant larvae have altered lipid levels. (A) Sterol levels are increased in *Lamp1*^*e879*^ mutants, an effect that is partially rescued by a duplication including *Lamp1* (Dp). Two-tailed T-tests. (B) Triacylglycerols (TAG) are unaffected. (C) Levels of larval DAGs and their indicated subclasses in *Lamp1*^*e879*^ mutants. Note the rescue by *Lamp1*^*Dp*^. One-way ANOVA (Tukey correction) significance levels are indicated below the X-axis. Unless indicated, changes are not significant (ns). *, P <0.05; **, P <0.01.

## Discussion

Our assessment of the function of Lamp1 using a marker of fluid phase endocytosis chased into lysosomes in living FB cells showed that Lamp1 localizes to late endosomes and lysosomes (Fig. 1), consistent with it being the homolog of the mammalian LAMP1/2 proteins, which is also supported by gene synteny of the *Lamp1* locus with vertebrates (Dr. L. Lescat and AJ; unpublished observation). Unexpectedly, *Lamp1* mutants are homozygous viable and show no developmental delay, distinct from mice lacking both, partially redundant *Lamp1/2* genes, which die around E15.5 [26]. We found a large expansion of the cellular acidic compartment under basal conditions and starvation in *Lamp1* mutants. Again, this is distinct from mouse *Lamp2* mutants and differentiated cardiomyocytes derived from hiPSC of Danon patients, in which no change in LTR staining was observed [31]. It is worth mentioning that lysosomal pH is driven by a difference in membrane potential between the cytoplasm and the lysosomal lumen (more positive), which could be altered in *Lamp1* mutants [60]. Interestingly, under basal conditions, the increase in acidic structures does not result in increased activity of lysosomal hydrolases, as we saw no increase in Cathepsin B or Acid Phosphatase activity (Fig. 3). Expanded acidic structures have been reported upon knockdown of *Drosophila Gga* (Golgi-localized, γ-adaptin ear containing, ARF binding protein) which is required for the transport of Cathepsins to lysosomes and thus for their proteolytic processing and activation. However, we found no indication of altered Cathepsin L processing in FB of *Lamp1* mutants (not shown), and the increase in LTR-positive puncta could thus represent an increase in late endosomes, as these organelles are acidified but lack hydrolytic activities [61,62].

*Lamp2* and *Lamp1/2* double mutant cells have a block in autophagic flux as shown by measurements of LC3-II levels [31] and of APGs using tandem GFP-RFP-LC3 reporters [25,31] and EM [22]. However, the mechanistic reason for this phenotype is unclear. It was suggested that LAMP2 is required for APG-lysosome fusion [25], or to promote fusion via simultaneous interaction with Atg14 on APGs and the lysosomal SNARE Vamp8 [31], therefore having a direct function in fusion. However, Hubert *et al*. [25] showed failure of APGs to recruit the SNAREs STX17 and SNAP29 required for fusion of APGs to lysosomes in *Lamp2* and *Lamp1/2* double mutant cells, which is not easily reconcilable with a direct function of LAMPs in MA. Additionally, non-functional vesicular transport and thus a potentially indirect effect has also been suggested as reason for defects in phagosome-lysosome fusion in *Lamp1/2* mutant MEFs [63]. Furthermore, the initial TEM analysis of *Lamp1/2* double mutant MEFs found no defect in the formation of APGs or their fusion with lysosomes, but rather a retardation in maturation of autolysosomes [26]. Our results show that basal MA flux and starvation induced MA are normal in *Drosophila Lamp1* mutant fat body (Fig. 4A-E). Consistently, we find normal recruitment of Snap29 to APGs (Fig. 4 F-H) and no accumulation of APGs in our TEM analyses, arguing against an essential direct function of LAMPs in MA. Alternatively, a more direct function of LAMPs in MA could have evolved after separation of vertebrates from arthropods. Furthermore, we cannot rule out that Lamp1 affects MA elicited by other forms of cellular stress. Importantly, we also show that, as in mammals [64,65], Lamp1 is dispensable for eMI and, since our eMI sensor is equivalent to the mammalian CMA sensor [53,57], our data functionally confirm that there is no evidence for CMA in *Drosophila*.

LAMP proteins have also been implicated in lipid metabolism [66]. LAMP1 and LAMP2 can bind cholesterol [58], and *Lamp1/2* double mutants accumulate cholesterol in the endolysosomal system [26,67]; thus LAMP proteins have been implicated in the handover of cholesterol to the NPC lysosomal cholesterol export system [58]. Consistent with those findings, Drosophila *Lamp1* mutant larvae also have increased sterols levels, although the increase is small compared with the effect observed in mice. Fruit flies are sterol auxotrophs and rely on dietary sterols for survival [68], and, consistently, both yeast-derived and plant-derived sterols were identified in our analysis (not shown).

Importantly, *Lamp1* mutant larvae also displayed a significant increase in DAGs. Unlike vertebrates, insects use DAGs as the main lipid species for transport of dietary lipids from the midgut to other tissues [59,69]. Lipids from the diet are digested by secreted lipases and esterases in the midgut, followed by absorption of fatty acids, sterols, and other molecules by enterocytes [59,69]. These cells then synthesize DAGs for export into the hemolymph, which transports lipids as lipoprotein complexes to other tissues [69]. Lipophorin (Lpp), a member of the ApoB family, is the main hemolymph lipid carrier [70]. The most abundant lipids in Lpp-containing lipoproteins are DAGs (70%), and phosphatidylethanolamine (20%), while sterols make up only 5% [70]. ApoLpp is synthesized in the FB and released to the hemolymph as high-density lipoproteins (HDL) that are loaded with phospholipids but lack DAGs. HDLs travel to the gut where they are loaded with DAGs with the help of another lipoprotein particle containing lipid transfer protein (LTP) and become low-density lipoproteins (LDL) [69,70]. LDLs are transported in the hemolymph for final use in organs [59,69].

Importantly, transported DAGs contain medium chain fatty acid, with a large proportion of C12 and C14 tails [70,71], corresponding to the species showing the most significant increase in *Lamp1* mutants (combined chain lengths of 24 and 28 carbons; Fig. 7). Thus, it is apparent that Lamp1 has a broader role in lipid assimilation and is not limited to sterol metabolism. However, the mechanism by which Lamp1 affects DAGs is less clear. Given Lamp1’s prevalent localization in late endosomes and lysosomes, it can be hypothesized that the protein has a role in endocytosis, and lack of Lamp1 could affect endocytosis-dependent lipid transport. For example, it was shown that Lpp loading partially depends on endocytosis, as the intermediary lipoprotein LTP needs to be internalized in midgut enterocytes for loading. In turn, LTP is necessary for loading of DAGs and sterols into Lpp for transport via the hemolymph [59,70]. Moreover, LTP RNAi results in large accumulation of medium chain DAGs in the gut [70] and defective LTP endocytosis in *Lamp1* mutants could explain the observed lipid phenotype.

Defects in phagocytosis have been previously reported in *LAMP1/2* mutant mice fibroblasts transfected with FcγIIA receptors [63]. In this case, phagosomes acquire Rab5 but not Rab7, and fail to fuse with lysosomes, indicating that endocytosis is impaired at a late step in the process. It was proposed that the endocytosis defect is associated with a failure of endosomes and lysosomes to move on microtubules [63]. Similarly, defective compensatory endocytosis was observed in *LAMP1/2* mutant fibroblast during plasma membrane repair [72]. The reduced ability of *LAMP1/2* mutant fibroblasts to carry out compensatory endocytosis also explains their increased resistance to *Trypanosoma cruzi* infection, as the parasite causes plasma membrane injuries and then hijacks the compensatory endocytosis process to enter the host cell [72,73]. However, in this case it was proposed that the role of LAMP2 in endocytosis may be indirect, as a decrease in cholesterol level in the plasma membrane associated with failure to recruit caveolin-1 could have led to a reduction in endocytosis [72]. Similarly, liver-stage *Plasmodium berghei* parasites have reduced growth in *LAMP1/2* mutant host cells because the parasite fails to recruit late endosomes and lysosomes to the parasitophorous vacuole (PV) [74]. The same phenotype was observed for *NPC* mutant host cells or when host cells were treated with the amphipathic steroid 3-b-[2-(diethylamine)eth-oxy]androst-5-en-17-one (U18666A). The common phenotype in the three cases is the accumulation of cholesterol in late endosome/lysosomes, suggesting that sequestration of cholesterol and potentially reduction of the available sterol pool leads to the defects in late endosome and lysosome recruitment to the PV, and/or reduced availability of sterols necessary for parasite growth [74]. These results further support the hypothesis that lack of LAMP proteins results in endocytosis defects, although the direct participation of LAMP proteins in specific membrane fusion events, or their indirect effect through changes in lipid composition of the membranes needs to be resolved. In any case, the disruption of endocytosis in *Lamp1* mutants could not completely block this process, since manipulation of *Drosophila* genes essential for endocytosis also disrupt lysosome biogenesis and result in an overall decrease in LTR-positive puncta [75], opposite to the phenotype observed for *Lamp1* mutants.

Our analysis indicated that *Drosophila* Lamp1 is necessary to maintain the normal cellular composition of acidic organelles, and for lipid transport. Our data showing that basal and starvation induced autophagy is not affected in *Lamp1* mutants, while sterol and DAG levels changed thus lend support to the previous suggestion that changes in sterol levels and the defects in autophagic flux observed in *Lamp1/2* mutant mice are independent phenotypes [67]. Moreover, the use of DAGs as the main species for lipid transport in insects rather than TAGs as in vertebrates allowed us to identify a broader effect for LAMP proteins in lipid metabolism without a confounding effect of lipid storage. These characteristics make *Drosophila* an ideal model to further dissect the roles of LAMPs in lipid metabolism and organelle traffic.

## Materials and Methods

### Fly strains and genetics

All fly stocks were maintained at 25□°C and reared on standard cornmeal media unless otherwise indicated. *PBac{RB}Lamp1*^*e00879*^ (here *Lamp1*^*e879*^) is a piggyBac insertion [76] in the first intron of Lamp1 (Fig. 1B) and is a RNA null allele (Fig. S1D). y1 M{nos-Cas9.P}ZH-2A w*(BL# 54591) was from the Bloomington stock center. Dp(2;3)*P6-B2*^*[Lamp1]*^ is a duplication including the *Lamp1* locus in VK31 on chromosome 3L (referred to here as Dp) that was obtained from Genetivision (Huston, TX). *UASp-GFP-mCherry Atg8a u2*.*6* (Chr. II; tandem tag) was from Dr. I. Nezis (U. Warwick, UK) and was recombined onto the *Lamp1*^*6*.*1*^ chromosome. *Lamp1*^*6*.*1*^; *r4-Gal4* and *Lamp1*^*11B*^; *r4-Gal4* used in the flux assays are based on *r4-Gal4* obtained from Dr. T. Neufeld (U. Minnesota)[77], who also provided *UAS-mCherry-Atg8a* lines [78]. *UAS-KFERQ-PAmCherry 3B* was as in [53]. See Table S1 for exact genotypes.

To measure pupation timing, flies were allowed to lay eggs overnight onto grape juice plates before removing hatched larvae by floating them on 20% sucrose for embryo synchronization. After verification of the absence of larvae, plates were incubated for 2 h at 25 °C and newly hatched larvae were floated off with 20% sucrose, collected, and placed at roughly 30 larvae/vial onto regular food (t= 25±1h to account for 24 h of embryogenesis). Pupating larvae were scored every 8 h.

### Molecular biology and transgenic flies

Transgenes were injected by Rainbow Transgenic flies (CA). pCFD4_Lamp_13 pCFD4_lamp1-30_4-4 were made by Gibson assembly (NEB # E5510S) using oligos Lamp1_crips_61_for, Lamp1_crisp_63_rev and Lamp1 exon 4_4, Lamp1 exon_1_30, respectively (Table S2) as described [38]. Transgenic lines were crossed with isogenized *nosCas9/FM6*; *Frt40/CyO*, allowing generation of *Lamp1*^*6*.*1*^ directly on a *FRT40* chromosome. Genotyping oligos are indicated in Table S1. *Lamp1*^*6*.*1*^ *FRT40* (*Lamp1*^*6*.*1*^ in text) contains frame shifts in exons 2 (5 bp deletion) and exon 3 (21 bp at expense of deletion of 8 bp, thus a net insertion of 13 bp) resulting in a predicted protein truncated after 90 aa of Lamp1 and an additional ectopic 32 residues (Figs. 1B and S1A). *Lamp1*^*11B*^ deletes all but the first 8 aa of the Lamp1 ORF including the stop codon (3067 bp, 1 bp insertion; 3066 bp net deletion; Figs. 1B and S1A).*Lamp1*^*6*.*1*^ *FRT40* was outcrossed 6 times against *w*^*1118*^ to lose an off-target lethal mutation. In both cases, non-targeted chromosomes were exchanged.

For qPCR, total RNA was extracted using TRIzol reagent (Thermo Fisher Scientific, Waltham, MA, USA) from ∼25-30 larvae or whole adult flies, DNase-treated (Turbo DNA-free kit; Ambion) and reverse transcribed using SuperScript III First Strand synthesis kit (Invitrogen). qPCR was performed in triplicates with an ABI prism 7300 Sequence Detection System (Thermo Fischer Scientific, Waltham, MA, USA) using Absolut qPCR with SYBR Green + Rox kit (Thermo Fischer Scientific; see Table S2 for primers). mRNA abundance was normalized with *RpL32* as housekeeping control using the ΔΔC_T_ method [79].

### Biochemistry and lipid analyses

For Western blot analyses, larvae were starved for 4 h on 20% sucrose and adults for 24 h with water saturated filter papers. 10 fly heads or 20 washed larvae were suspended in 50 µl or 200 µl 1x Laemmli buffer with 2% SDS (2% SDS, 10% glycerol, 5% 2-mercaptoethanol, 0.002% bromophenol blue and 62.5 mM Tris HCl, pH 6.8), respectively, and boiled for 5 min at 95 °C. After homogenization with a motor pestle, lysates were boiled again and centrifuged twice for 10 min at 20,000 g at room temperature. After each centrifugation, the 90% of the liquid phase was removed avoiding lipids floating on top. Upon separation of the proteins by SDS-PAGE, proteins were transferred to nitrocellulose membranes and probed with NY2403 anti Lamp1 (see below) at 1:5000 and anti αTubulin (Sigma, T5168; 1:10,000) as loading control.

For deglycosylation assays, 40 adult heads of indicated genotypes were lysed in 200µl RIPA buffer (150 mM NaCl, 5 mM EDTA pH 8.0, 20 mM Tris pH 8.0, 1% NP40, 0.25 % Deoxycholate) by homogenization with a motor pestle. After clearing at 10,000 g for 10 min at 4°C, 70 µg protein (as determined with a Lowry assay) was combined with 3 µl denaturing buffer (5% SDS, 400 mM DTT), heated to 95°C for 10 min prior to addition of protease inhibitors. 2 µl Endo H (NEB P0702S) and 3 µl of G3 reaction buffer (NEB, MA) and 2 µl PNGase F (NEB P0704S), 3µl 10% NP40, and 3µl G2 reaction buffer were added, respectively. After incubation for 18 h at 37°C, samples were boiled in Laemmli buffer and processed for Western blot analysis.

To measure Acid Phosphatase activity, a fraction containing lysosomes, mitochondria, and cytoplasm was prepared by centrifugation from 15 to 30 3^rd^ instar larvae as described [46], with the addition of cOmplete, Mini, EDTA-free Protease Inhibitor Cocktail (Sigma) in the extraction buffer. The presence of lysosomes in this fraction was confirmed by Lysotracker staining and observation under fluorescence microscope. Protein concentration was determined using the Pierce BCA protein assay kit (Thermo Fisher Scientific). Acid phosphatase activity was determined fluorometrically using 4-methylumbelliferyl phosphate as substrate [80].

For lipid quantification, ten 3^rd^ instar larvae were weighed and homogenized with 350 µl of hot methanol (60°C), spiked with 25 μg of ribitol and 25 μg of nonadecanoic acid as internal standards. The mixture was immediately incubated at 60°C for 10 min and sonicated for 10 min. Samples were then extracted with 350 µl of chloroform and 300 µl of water and centrifuged for 7 minutes at 14,000 g. The lower, non-polar fraction was transferred to new vials and dried in a Speed-Vac concentrator. Non-targeted metabolite analysis was used to quantify sterols and diacylglycerols (DAG). The samples were methoximylated and silylated as described[81]. One microliter of the derivatized samples was injected into an Agilent 6890 GC interfaced to an Agilent 5973 quadrupole MS with a HP-5ms (5%-Phenyl)-methylpolysiloxane column (30 m × 0.25 mm × 0.25 μm, Agilent) in splitless mode. The temperature was programmed from 70 to 320°C at 5°C/min with helium flow rate at 1.0 mL/min and inlet temperature at 280°C. EI-MS ionization energy was set to 70 eV and the interface temperature was 280°C. The GC-MS data files were deconvoluted and searched against an in-house MS-library and the NIST 14 Mass Spectral Library using NIST AMDIS software [82]. Triacylglycerols (TAG) were analyzed with an Agilent Technologies 1200 Series four solvent gradient capable HPLC coupled to Agilent 1200 series evaporative Light scattering detector (ELSD) and a Supelco Ascentis-Si (25cm x2.1 mm x5 um) column. Elution was performed using mobile phases containing A: hexane with 1% isopropanol with 0.4% acetic acid added, and B: 100% isopropanol. The solvent gradient used was: 0-5 min 100% solvent A, 5-10 min gradient to 95% solvent A, 10-15 min 95% solvent A, 15-20 min 100% solvent A; with a flow rate of 0.8 ml/min. ELSD: Nitrogen flow was 2.1 with 110°C source temperature. A mixture of TAGs (2 mg/ml) was used as quantification standard. All lipids were normalized using the dry weight after extraction.

### Autophagy, Lysotracker, Magic Red, and Dextran assays

Dextran uptake assays to label lysosomes was performed as described [35,36]. Briefly, fed or starved larvae were washed, inverted and incubated with fluoro-Ruby-Dextran (1:100 of 100 mg/ml stock; 10,000 MW; Invitrogen CA) in Graces medium for 10 min, followed by 6 washes and a 90 min chase to allow the Dextran to be transported to lysosomes. Fixation and staining was as described below. Co-localization was calculated using Pearson’s correlation coefficient, turning one channel by 180° as control [37].

MA flux was determined using a tandem GFP-RFP-Atg8a reporter as described [50]. Briefly, well-fed mid 3^rd^ instar larvae of appropriate genotypes were washed with H_2_O and incubated for 3-4 h in 35 mm petri dishes with 3 Whatman filter paper soaked in 800 µl 20% sucrose supplemented with heat-inactivate yeast (fed) or 20% sucrose only (for starvation) [48]. After dissection and fixation, FB lobes of approximately 5 larvae were mounted in 20 µl DAPI fluoromount-G (Southern Biotech 0100-20) per slide and imaged on an ApoTome.2 system using an Axiovert 200 equipped with a 63x 1.4 NA oil lens (Carl-Zeiss, Oberkochen, Germany). Quantification was done using the atgCOUNTER script in Fiji to threshold and exclude nuclear signal [50].

eMI activity was determined as described[53] after recombining *r4-Gal4* with *UAS-KFERQ-PAmCherry 3B* (once in *Lamp1*^*6*.*1*^ background, the strain was backcrossed to homozygous *Lamp1*^*6*.*1*^ flies to consistently have only one copy of the reporter construct). FB was imaged as above.

For Lysotracker staining, 3^rd^ instar larvae were starved in PBS for 4 h (Fig. 2) or 20% sucrose (Fig. S2). Larval or adult FB Larval and adult flies fat body tissues were dissected in 1X PBS, incubated for 3 to 5 min in 100 μM Lysotracker-Red DND-99 (Life Technologies, Carlsbad, CA, USA)[49] and 1 μM Höchst 33342 (Thermo Fisher Scientific Inc, Rockford, IL, USA) in PBS and immediately imaged using a Zeiss Imager (larval FB) or an Olympus BX51WI (adult tissues) using identical conditions across genotypes. FB cells that contained any LTR-positive puncta in adult flies or FB cells that contained more than 5 LTR-positive puncta in larvae were scored as positive.

For Magic red staining (Cathepsin-B Assay Kit; Immunochemistry Technologies LLC, Bloomington, MN, USA), larvae were starved for 4 h in 20% sucrose (supplemented with heat inactivated yeast for fed controls), washed and dissected in Graces medium (Invitrogen, CA; without serum in case of starved larvae). Larvae were inverted in the corresponding Graces medium and stained for 10 min in the same medium supplemented with Magic red substrate (total dilution 1:150) and DAPI (14.3 M stock at 1:2000). After 2 rinses in corresponding Graces medium, FB was dissected on glass slides and immediately imaged on a Zeiss ApoTome with a 40x/1.3NA oil DIC lens.

### Immunohistochemistry and EM analyses

Lamp1 antibodies were generated against KLH coupled C-terminal peptide (CARRRSTSRGYMSF) in rabbits (#NY2403) by Covance (Denver, PA; animal protocol # 20170902) and affinity purified for use in immunohistochemistry (1:400-1:1000). Staining of FB was done as described [53,83]. Rabbit anti Snap29 [51] was used at 1:1000. Secondary antibody Alexa488 anti-rabbit (A11034; Invitrogen, CA) was used at 1:300.

For TEM, larvae were treated and processed as above, and, after inversion, fixed with 2.0% paraformaldehyde, 2.5% glutaraldehyde in 0.1 M sodium cacodylate buffer, postfixed with 1% osmium tetroxide followed by 2% uranyl acetate, dehydrated through a graded series of ethanol and embedded in LX112 resin (LADD Research Industries, Burlington VT). Ultrathin sections were cut on a Leica Ultracut UC7, stained with uranyl acetate followed by lead citrate and viewed on a JEOL 1200EX transmission electron microscope at 80kv. Morphometric quantification was done by outlining indicated structures in Fiji/ImageJ and tabulating their area [84] on blindly and randomly selected fields of view (excluding only nuclei). APGLs were identified as electron dense (dark) structures with very well or less-well (likely more degrading already) defined content, in contrast to ALs that were less electron dense (more similar to surrounding cytoplasm) and contained ill-defined content (advanced degradation). APGLs and ALs are likely similar to AVds in e.g. [18]. Lysosomes (L) are very electron dense without halo (distinguishing them from peroxisomes) and small. Structure density (are per field of view of 100 µm^2^) or size distribution was analyzed using the Mann-Whitney and Kolmogorov-Smirnov tests.

### Statistical analysis

Statistical tests were done with GraphPad Prism (Versions 8&9; GraphPad Software, La Jolla, CA) using indicated tests and corrections. Unless noted, graphs represent means with standard error of means (SEM). ‘n=‘ indicates fields of view unless indicated. As larvae are bilaterally symmetric and have two main FB lobes, we cannot exclude that two fragments may have originally belonged to the same lobe in the preparation. It is therefore a reasonable estimate that the number of animals is minimally about n/2 and maximally equal to n.

## Acknowledgements

We thank Drs. T. Neufeld, I. Nezis, G. Juhasz, T. Vaccari, and the Bloomington *Drosophila* Stock centers for kindly sharing fly strains. We are grateful to Dr. T. Vaccari for antibodies and to Drs. Mimi Kim (Division of Biostatistics, Einstein) and Ana Maria Cuervo for advice on the analyses of TEM images. We thank Dr. L. Ambrosio for help with fly genetics, the W. M. Keck Metabolomics Research Laboratory at Iowa State University for help with lipidomics analyses, and Drs. J. Secombe and A. Melendez for comments on this manuscript and the Einstein Analytical Imaging Facility (NIH P30CA013330) for support. This work was supported by AHA postdoctoral fellowship 18POST34030231 (to A.M), NIH/NIGMS grant GM119160 (to A.J.), and grants of the Roy J. Carver Charitable Foundation (Muscatine, Iowa) and NSF grant MCB– 1714996 (to G.C.M.). A.R and N.C. were further supported by Fulbright fellowships.

## Abbreviations

aa: amino acid
AL: autolysosome
APG: autophagosome
APGL: autophagolysosome
AV: autophagic vacuole (i.e. APG and APGL/AL)
AVi: early/initial autophagic vacuoles
AVd: late/degradative autophagic vacuoles
*Atg*: autophagy-related
CMA: chaperone mediated autophagy
DAG: diacylglycerol
eMI: endosomal microautophagy
ESCRT: endosomal sorting complexes required for transport
FB: fat body
HDL: high density lipoprotein
LAMP: lysosome associated membrane protein
LD: lipid droplet
LDL: low density lipoprotein
Lpp: lipophorin
Ltp: Lipid transfer protein
LTR: Lysotracker
MA: macroautophagy
MEF: mouse embryonic fibroblast
mTORC: mechanistic target of rapamycin complex
PV: parasitophorous vacuole
SNARE: soluble N-ethylmaleimide sensitive factor attachment protein receptor
Snap: synaptosomal-associated protein
st: starved
TAG: triacylglycerol
TEM: transmission electron microscopy
TFEB: transcription factor EB
TM: transmembrane domain
tub: tubulin
UTR: untranslated region.

## Figure legends

**Figure S1.**(A) Sequence of WT (upper line) and indicated *Lamp1* Crispr alleles (respective lower lines). Net deletions are indicated. In *Lamp1*^*6*.*1*^, the first of the two frame shifts results in addition of 32 non-Lamp1 aa after aa 90 (well upstream of the transmembrane domain). gRNA target sequences are indicated in blue, start/stop codons in red, and inserted bases are in green. (B) *Lamp1*^*e879*^ is an RNA null allele. Note that the duplication Dp(2;3)*P6-B2*^*[Lamp1]*^ uncovering Lamp1 (Dp) rescues expression to normal levels. *RpL32* was used as positive control. (C, D) Western blots of fed (F) and 4 h starved (ST) adult head (C) and 3^rd^ instar larval (D) lysates show that Lamp1 migrates as a major form of ∼45 kDa with several minor forms up to ∼60 kDa (magenta arrowheads; compare WT with *Lamp1*^*6*.*1*^ and *Lamp1*^*11B*^ lanes for specificity). Blue arrowheads indicate predicted molecular mass. αTubulin was used as loading control shown in lower panels. (E-H) Compared to fed and starved fat body of *w*^*1118*^ 3^rd^ instar FB (E,G), *Lamp1*^*6*.*1*^ mutants (F,H) lack Lamp1 staining (green), demonstrating specificity of the antibody also for histochemistry. Nuclei in blue. Scale bar: 10µm.

**Figure S2.**Increased LTR staining in 3^rd^ instar FB of *Lamp1* mutants. (A, D) *w*^*1118*^ control. (B,E) *Lamp1*^*6*.*1*^ and (C,F) *Lamp1*^*11B*^ mutants. A-B: fed, D-F starved conditions. LTR in red, nuclei in blue; scale bar: 20 µm.

**Figure S3.**TEM analyses of density (area per field of view) of lipid droplets (A) and glycogen (B) in fed and starved WT and *Lamp1*^*6*.*1*^ mutant larvae. Mann-Whitney tests; ns: not significant.

