## Supplementary figures and images for "Lamp1 mediates lipid transport, but is dispensable for autophagy in *Drosophila*"

### Supplementary Figure 1

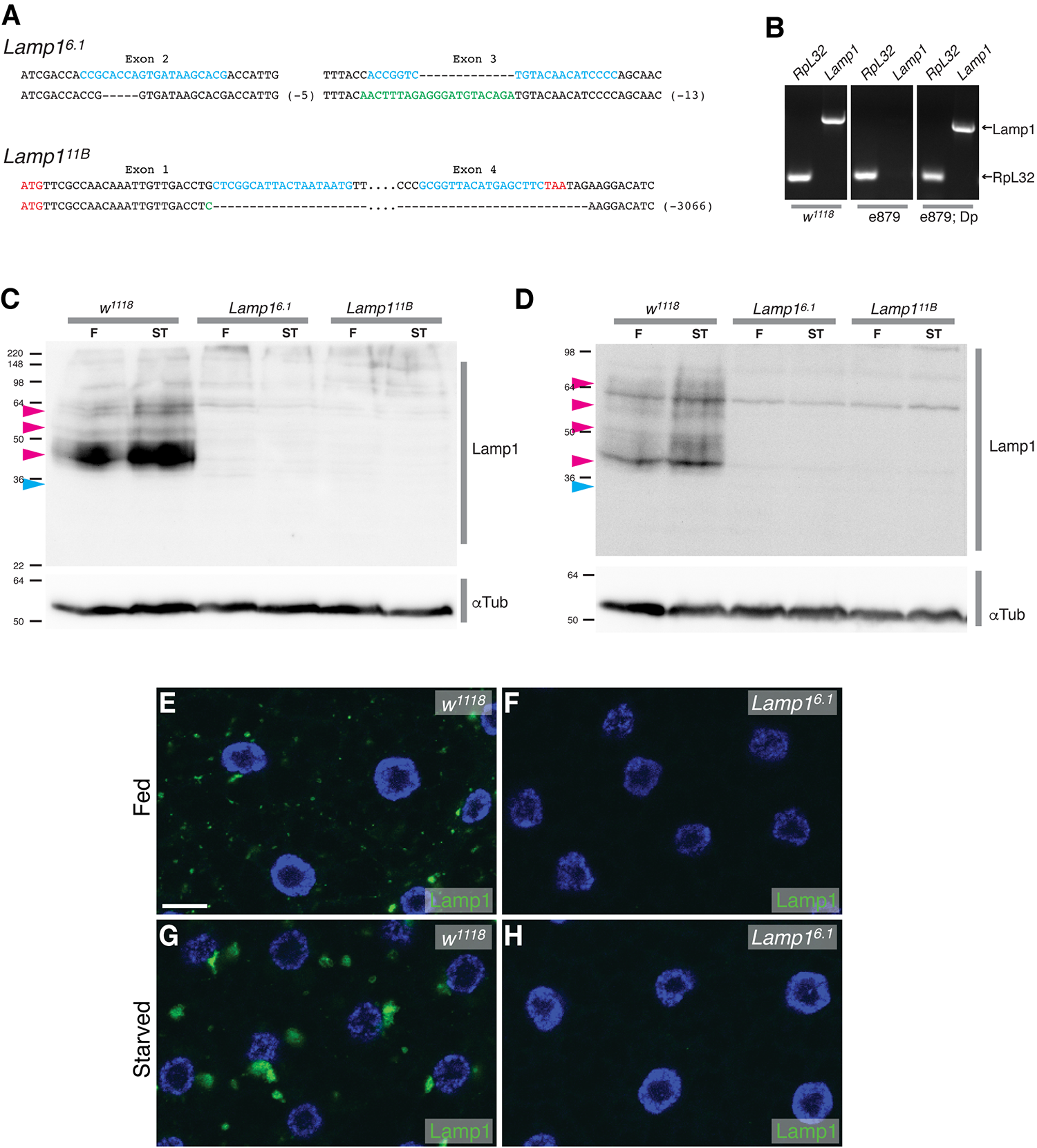

### Supplementary Figure 2

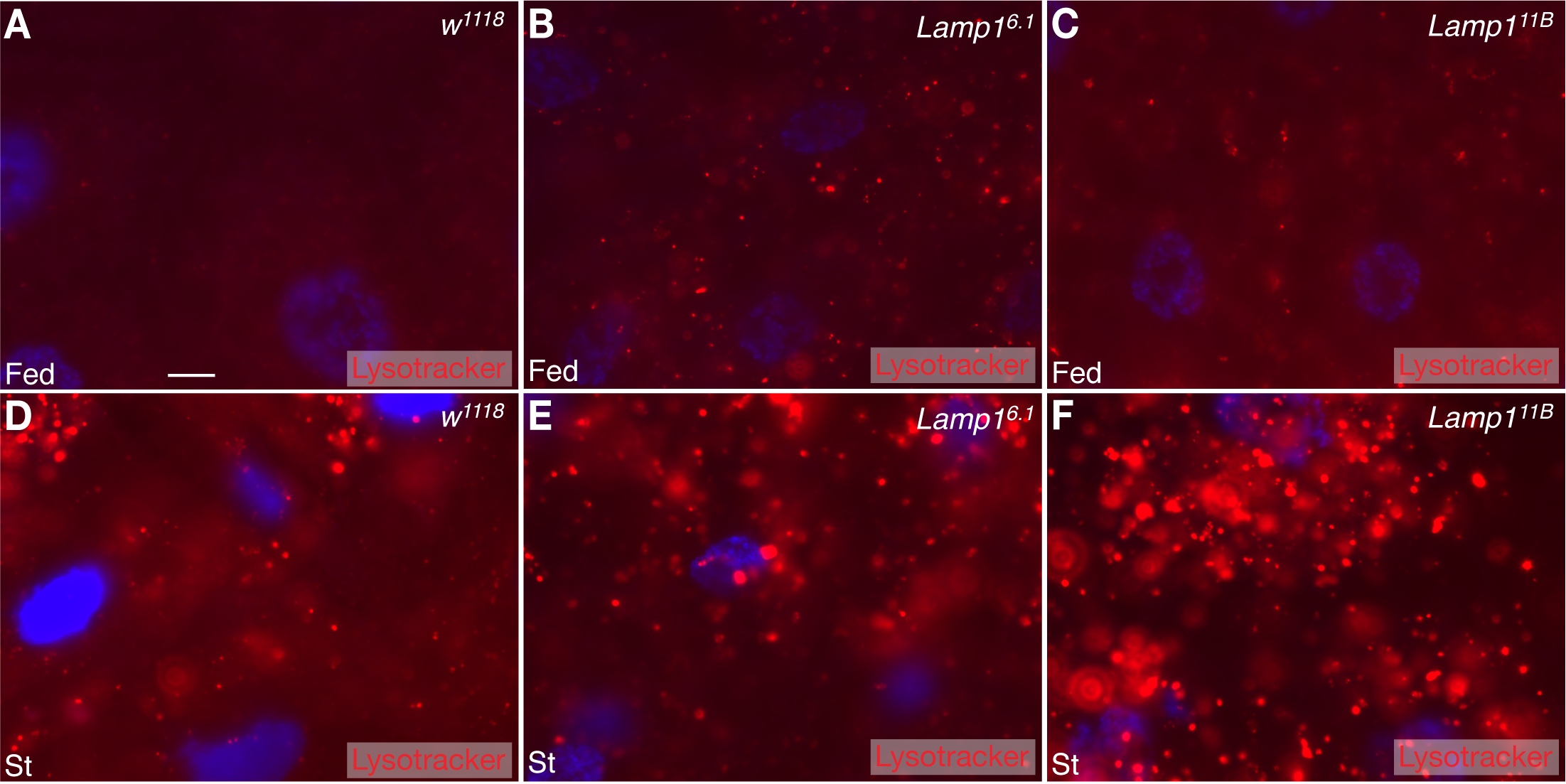

### Supplementary Figure 3

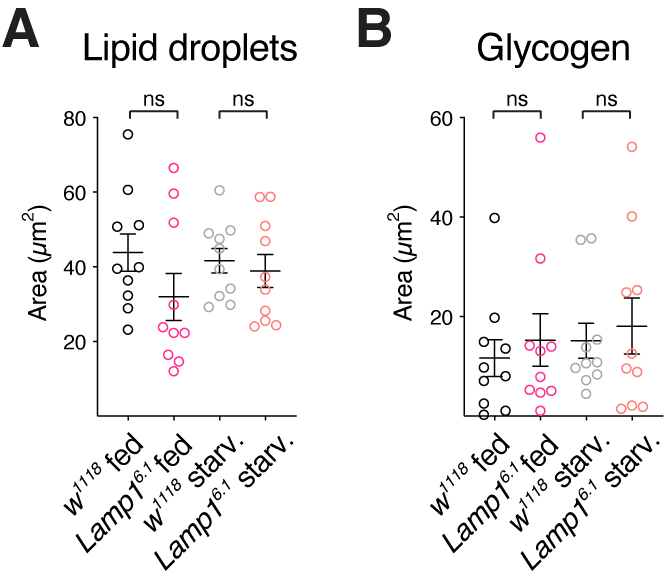
