## Supplementary Table 1 for "Lamp1 mediates lipid transport, but is dispensable for autophagy in *Drosophila*"

**Table S2:** Genotypes

**Figure 1:**

**C:**	*w^1118^*

*w^1118^*; *Lamp1^6.1^ FRT40*/ *Lamp1^6.1^ FRT40*

**D, E:**	*w^1118^*

**G:**	*w^1118^*

*w^1118^*; *Lamp1^6.1^ FRT40*/ *Lamp1^6.1^ FRT40*

**Figure 2:**

**A, E, G:** *w^1118^*

**B, F, H:**	*w^1118^; Lamp1^e879^*/ *Lamp1^e879^*

***C:***	*w^1118^*; *Lamp1^11B^*/ *Lamp1^11B^*

***D:***	*w^1118^; Lamp1^e879^*/ *Lamp1^e879^; Dp(2;*3*)P6-B2^[Lamp1]^ Dp(2;*3*)P6-B2^[Lamp1]^*

**Figure 3:**

**A, D:**	*w^1118^*

**B, E:**	*w^1118^*; *Lamp1^6.1^ FRT40*/ *Lamp1^6.1^ FRT40*

**C, F:**	*w^1118^*; *Lamp1^11B^*/ *Lamp1^11B^*

**Figure 4:**

**A, C, F:**	*w^1118^*; *Lamp1^6.1^ FRT40*/ *+*

**B, D, G:** *w^1118^*; *Lamp1^6.1^ FRT40*/ *Lamp1^11B^*

**I:**	*w^1118^*

*w^1118^; Lamp1^e879^*/ *Lamp1^e879^*

**Figure 5:**

**A, C:**	*w^1118^*; *Lamp1^6.1^ FRT40*/ *+*

**B, C:**	*w^1118^*; *Lamp1^6.1^ FRT40*/ *Lamp1^6.1^ FRT40*

**Figure 6:**

**A, B, C, E:** *w^1118^*; *Lamp1^6.1^ FRT40*/ *Lamp1^6.1^ FRT40*

**D:**	*w^1118^*

**Figure 7:**

**A-C:**	*w^1118^*

*w^1118^; Lamp1^e879^*/ *Lamp1^e879^*

*w^1118^; Lamp1^e879^*/ *Lamp1^e879^; Dp(2;*3*)P6-B2^[Lamp1]^ Dp(2;*3*)P6-B2^[Lamp1]^*

**Figure S1:**

**B, C:**	*w^1118^*

*w^1118^*; *Lamp1^6.1^ FRT40*/ *Lamp1^6.1^ FRT40*

*w^1118^*; *Lamp1^11B^*/ *Lamp1^11B^*

**D:**	*w^1118^*

*w^1118^; Lamp1^e879^*/ *Lamp1^e879^*

*w^1118^; Lamp1^e879^*/ *Lamp1^e879^; Dp(2;*3*)P6-B2^[Lamp1]^/ Dp(2;*3*)P6-B2^[Lamp1]^*

**E, G:**	*w^1118^*

**F, H:**	*w^1118^*; *Lamp1^6.1^ FRT40*/ *Lamp1^6.1^ FRT40*

**Figure S2:**

**A, D:**	*w^1118^*

**B, E:**	*w^1118^*; *Lamp1^6.1^ FRT40*/ *Lamp1^6.1^ FRT40*

**C, F:**	*w^1118^*; *Lamp1^11B^*/ *Lamp1^11B^*

**Figure S3:**

**A, B:**	*w^1118^*

*w^1118^*; *Lamp1^6.1^ FRT40*/ *Lamp1^6.1^ FRT40*
