## Supplementary Table 2 for "Lamp1 mediates lipid transport, but is dispensable for autophagy in *Drosophila*"

**Table S1 Oligonucleotides**

**Crispr gRNA cloning:**

Lamp1_crips_61_for (Cr 8; into U6-1 site)

TATATAGGAAAGATATCCGGGTGAACTTCGGGGATGTTGTACAGACCGGGTTTTAGAGCTAGAAATAGCAAG

Lamp1_crisp_63_rev (Cr4; into U6-3 site)

ATTTTAACTTGCTATTTCTAGCTCTAAAACCCGCACCAGTGATAAGCACGCGACGTTAAATTGAAAATAGGTC

Lamp1 exon 4_4 (into U6-1 site)

TATATAGGAAAGATATCCGGGTGAACTTCGTTAGAAGCTCATGTAACCGCGTTTTAGAGCTAGAAATAGCAAG

Lamp1 exon_1_30 (into U6-3 site)

ATTTTAACTTGCTATTTCTAGCTCTAAAACGCTCGGCATTACTAATAATGCGACGTTAAATTGAAAATAGGTC

**Genotyping primers:**

Lamp1Cr4_test_for: CTCAACAATCTCAACGTCTTCG

Lamp1Cr4_test_rev: AGGTAAAATTAAGTTGTGCCGC

Lamp1Cr4_rev_intronic: CCTCACCAATGCCGTAATTC (for genotyping rescued strains)

Lamp1Cr8_test_for: TAATGGGCAAACGCTTTTTATT

Lamp1Cr8_test_rev: ATTGCTTTGATGTTTCTGGTCC

Lamp1Cr1_30_test_for: ATTACGGTTCCATATGGGCTAA

Lamp1Cr1_30_test_rev: GACGCATTTCATTGCTGTTTTA

Lamp1Cr4_4_test_for: AGTTAATTGGTTGGATCAGCGT

Lamp1Cr4_4_test_rev: TTTGCTGTCCATACTCCTTTGA

**qPCR**

RpL32_for: AAGATCGTGAAGAAGCGCAC

RpL32_rev: CGTAACCGATGTTGGGCATC

Lamp1_RT_for: CTAGCTCAGTCGAGACAAAAACG

Lamp1_RT_rev: AGGCAGCAAAGCCAGATGAA

Atg5_for: ATCTGGGAGGGCCAGATAGG

Atg5_rev: TAGCTCCTTGGAGTTGAGCTTG

Atg8_for: CCTGTACCAGGAACATCACGA

Atg8_rev: GGGTAGGACACAAAGCAGAGT

Amyrel_for: GATCTAGAGTACATCTACAGCAGCC

Amyrel_rev: ACTTGTAGTTCAGCACGGCA
